# The influence of sleep disruption on learning and memory in fish

**DOI:** 10.1101/2024.08.28.610197

**Authors:** W. Sowersby, T. Kobayahsi, S. Awata, S. Sogawa, M. Kohda

## Abstract

Sleep is ubiquitous across animal taxa. Strong evolutionary pressures have conserved sleep over the evolutionary history of animals, yet our understanding of the functions of sleep still largely derives from mammals and select laboratory models. Sleep is considered to play an important role in mental processes, including learning and memory consolidation, but how widespread this relationship occurs across taxa remains unclear. Here, we test the impact of sleep disruption on the ability of the cleaner fish *(Labroides dimidiatus)* to both learn and then remember a novel cognitive task. We found a significant negative relationship between sleep disruption and the ability to learn a food reward choice system. Specifically, we found that fish in a disturbed sleep treatment took significantly longer and made more incorrect decisions when finding the food reward, compared to individuals in a non-disturbed/normal sleep treatment. In contrast, the differences between the two treatment groups were non-significant when fish where tasked with remembering the food reward several days later. Our results demonstrate a negative impact of sleep disruption on performance in a cognitive challenging task and that the effects were strongest when fish were first exposed to the challenge. Importantly, we show that the association between sleep and mental processes, such as learning, may be widespread across vertebrate taxa and potentially have an early origin in the evolutionary history of animals.

## Introduction

Sleep is a ubiquitous trait that has been conserved across animal taxa, with positive links to immune function, neurogenesis, wellbeing, and cognitive performance (Tononi & Cirelli 2012; Czeisler 2015; Elvsåshagen et al. 2015). There are also costs associated with sleeping, such as potentially being more vulnerable to predators and reducing the time available to secure food and mates (Lima et al. 2005; Joiner 2016; Tworkowski & Lesku 2019). Nevertheless, there is currently no evidence to suggest that animals can completely renounce sleep without detrimental consequences, including death (Van Dongen et al. 2003; Cirelli & Tononi, 2008). Indeed, a commonly observed response to sleep disruption, is longer and deeper sleep, indicating that sleep is maintained homeostatically (Tobler 2011; Tobler and Borbély 1985; Nath et al. 2017; Yokogawa et al. 2007). Despite seemingly strong selective pressure to conserve sleep across animal evolutionary history, how sleep has coevolved with other traits and processes is still not often considered in evolutionary research (Cirelli and Tononi 2008; Mignot 2008; Roth, Rattenborg, and Pravosudov 2010; Rattenborg et al. 2017).

One exciting but still controversial proposed function of sleep is the important role it plays in brain processes. For example, compelling evidence points to a strong positive link between sleep, learning and memory consolidation (Huber et al. 2004; Walker and Stickgold 2004b; Peigneux et al. 2001; Smith 2001; Walker and Stickgold 2006; Anafi, Kayser, and Raizen 2019; Siegel 2001; but see Vertes 2004; Rattenborg et al. 2017), neural maintenance (Kavanau 1996; Tononi and Cirelli 2006) and cognitive ability (Scullin and Bliwise 2015; Wagner et al. 2004). In animals, learning and memory are critical cognitive functions for a range of tasks, including foraging or food-caching (Shettleworth 1995; Sherry and Hoshooley 2010), navigation and territory maintenance (Shettleworth 1998) and for recalling positive and negative social interactions (Mateo and Johnston 2000). The link between sleep and memory consolidation has been demonstrated in humans (Born, Rasch, and Gais 2006; Diekelmann and Born 2010; Huber et al. 2004) and other mammals (Capellini et al. 2009; Smith and Kelly 1988; Ribeiro et al. 2004; Walker and Stickgold 2004b) and to a lesser extent in birds (Jackson et al. 2008; Derégnaucourt et al. 2005). To date, research investigating this relationship has largely been restricted to these few taxa (Keene and Duboue 2018), despite some evidence indicating the importance of sleep for memory formation in some invertebrates (Beyaert, Greggers, and Menzel 2012; Krishnan, Noakes, and Lyons 2016) and fishes (Pinheiro-da-Silva, Tran, and Luchiari 2018; Rawashdeh et al. 2007).

The function and in places the structure of the vertebrate brain, especially the forebrain, appear to have been largely conserved throughout the evolution of vertebrate taxa (Vargas et al. 2006; Xie & Dorsky 2017). For example, the mammalian and avian hippocampus, along with the homologous pallial areas of the reptile and fish brain, share a central role in supporting and encoding spatial information (Rodriguez et al. 2002; Portavella et al. 2002). Vargas et al. (2006) demonstrated that lesions to the lateral pallium of goldfish (*Carassius auratus*) can impair the encoding of geometric spatial information, comparable to the effect of lesions on the hippocampus in mammals and birds. Interestingly, comparable patterns of sleep have also been observed across these taxonomic groups, including in fishes, which display both characteristic sleep behaviours (in cartilaginous fishes: Kelly et al. 2020, 2022) and deep slow-wave and rapid eye like movements (REM; in bony fishes: Leung et al. 2019; M. Kohda et al. unpublished data). Evidence suggesting an association between learning and active memory consolidation with sleep in non-mammalian species would imply that sleep has played an important role in these mental processes across vertebrate evolutionary history (Vorster and Born 2015). Any link between sleep, learning and memory in fishes or other non-mammalian taxa is however not well established (Pinheiro-da-Silva, Tran, and Luchiari 2018; Rawashdeh et al. 2007) and this large and diverse group of vertebrates has largely been neglected from studies on sleep (Campbell and Tobler 1984).

Here, we investigate the impact of sleep disruption on learning ability and subsequent memory consolidation in the bluestreak cleaner wrasse (cleaner fish; *Labroides dimidiatus*). This fish species has a wide distribution on coral reefs across the Indo-Pacific, where in a mutualistic relationship, it eats parasites and dead tissue off “client” species. Hence, this highly social fish, regularly engages in cognitively challenging tasks, such as recalling both favourable and antagonistic interactions with both conspecifics and numerous client individuals from a range of different species. Cleaner fish are highly amenable to captive experimental conditions and have demonstrated several complex cognitive abilities, some which may be impacted by sleep quality (e.g. Bshary & Brown 2014; Kohda et al. 2019, 2023). We predict that when subjected to interrupted sleep schedules, cleaner fish will demonstrate poorer performance in novel cognitively challenging tasks, in comparison to fish that have experienced uninterrupted, normal sleep regimes. We also predict that cleaner fish subjected to disruptions in normal sleep regimes will take longer to rouse each morning and react slower to the presence of food, compared to fish that have experienced uninterrupted sleep.

## Methods

### Animal husbandry

Cleaner fish (n=12) were purchased from reputable tropical fish specialists and were individually housed (aquaria 40L) under laboratory conditions (14:10 hr light regime; 27 degrees C) at the Department of Biology, Osaka Metropolitan University, Japan. All aquaria were furnished with coral sand substrate and a small opaque piece of PVC pipe to provide shelter. Water changes and water parameter checks (including salinity) were conducted regularly. Fish were fed daily with commercially bought shrimp (Pandalidae), mashed into a paste and suspended in the tank on a grey plastic plate. A pilot trial of the experimental set-up took place in early 2020, with the main experiment then conducted between September and October 2023. All experiments adhered to the animal ethics guidelines of Osaka Metropolitan University.

### Sleep Treatment

Fish were moved into their treatment tanks for a 48-hour acclimation period prior to the start of the experiment. Treatment tanks were identical to housing aquaria, except treatment tanks were divided into two by a clear plastic barrier with a door and had a lamp overhead (Figure 1). Fish were placed into the back half of the tank and were provided with food in the front half of the tank, once per day. Specifically, shrimp mashed into a paste was put on a grey plastic square which was hung into the front half of the tank. When the door in the partition was raised the fish could enter the front half of the tank and became accustomed to accessing that half of the tank via the door to feed. After three hours the grey plastic square was removed, and fish were gently ushered back into the second half of the tank if they had not already returned voluntarily.

**Figure 1.**
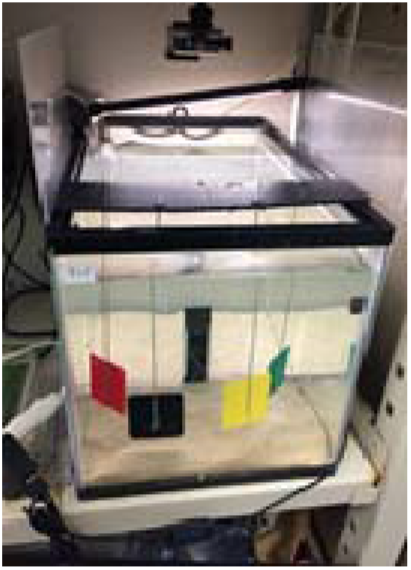
Set-up for the food choice reward experiment.

After the acclimation period, half of the fish (n = 6) were assigned to the interrupted sleep treatment and the remaining half to the non-interrupted sleep treatment (n = 6). Fish in the non-interrupted sleep treatment were kept in complete darkness for a period of 10 hours (22:00-8:00) each night, a period which corresponds to the typical night length in the tropical regions where these fish naturally occur. In contrast, fish in the interrupted sleep treatment were exposed to a light that came on above the tank every two hours, for a period of one hour (controlled by a timer), meaning that fish in this treatment were exposed to less periods of darkness per night, broken into two-hour blocks. Using light to supress sleep has successfully been used in zebrafish research (Yokogawa et al. 2007). Moreover, the opaque pipe shelter was removed in the interrupted sleep treatment and replaced with a transparent plastic pipe of identical size, which would provide the fish with a secure place to shelter, but not offer refuge from the light above.

To prevent fish in the non-interrupted sleep treatment from being disturbed by light during the night, fish in the sleep interrupted treatment were isolated at night via a black-out sheet. Camcorders (SONY Handycam HDR-CX480) positioned in front of each tank (n = 4) were quietly switched on 10 minutes prior to the morning light coming on (at 8:00). The room was blacked out to prevent natural light entering, and researchers entered using a dim red light. Fish were recorded for ∼20 minutes after the morning light automatically switched on at 8.00am, to capture the time it took each fish to wake up (i.e. when they left the shelter and started swimming), and their initial activity behaviour (i.e. time spent moving and distance travelled). This sleep treatment occurred every night for four nights during the learning phase of the experiment. During the subsequent rest and memory phase of this experiment all fish were exposed to darkness for the normal 10-hour period and were provided with an opaque tube for shelter. A subset of the fish were recorded during the night, confirming that their sleep was disturbed by the light treatment.

### Food reward choice: Learning phase

We presented fish with a food reward choice at ∼9.00am each morning, for three days, beginning from the first morning after the night of the first sleep treatment. Each fish was presented with a row of four plastic squares, each a different colour (red, green, black and yellow) suspended in a row in the front half of the tank (Figure 1). Shrimp paste had been placed on the back of only the red square, so fish were tasked with finding the food on the back of this square. Each morning for the three days, fish were presented with the same four coloured squares, but with the order of the squares in the row different each time. Once the squares were in place the door separating the front and back halves of the tank was lifted and we recorded the following, 1). time to exit the door, 2). number of incorrect choices, 3). time to find food on the back of the red square. We considered an incorrect choice to be any deliberate touching of a square with the mouth or head that was not on the backside of the red square. After three mornings, fish in the sleep interrupted treatment were exposed to a final night of disrupted sleep before all fish entered a six-day rest period.

### Food reward choice: Memory phase

After the rest period, fish were again exposed to the same food reward choice experiment for another three days. Like the rest period, no fish were exposed to the non-interrupted sleep treatment. This experiment tested the ability of the cleaner wrasse to recall the correct coloured square to locate the food reward. As previously, we recorded, 1) time to exit the door, 2) number of incorrect choices, 3) time to find food on the back of the red square. After the third day each fish was weighed and photographed and placed back in its home tank.

### Statistical Analysis

We performed separated repeated measures ANOVA to test the effect of sleep disruption on wake time, time to exit door, number of incorrect choices and time to correct choice. In each model we included the variables sleep treatment (disrupted or non-disrupted) and day (day 1, 2 or 3) with experimental tank added as a random effect. All statistical analyses were conducted in R v4.2.1 (R Core Team, 2022).

## Results

### Impact of sleep disruption on ability to learn cognitive task

In contrast to our first prediction, we did not observe any significant difference in time to first movement between fish in the disturbed and non-disturbed sleep treatments (rousing time; df=1, F=0.018, p=0.896) across the four mornings after sleep disturbance. On average, fish in the disturbed sleep treatment first moved from their shelter after 197 seconds and non-disturbed treatment fish 220 seconds once the light automatically turned on at 8.00am.

Non-disturbed sleep fish did exit the door quicker than disturbed fish, however this difference was not significant (df=1, F=1.552, p=0.241).

In line with our second prediction, we did find a significant negative effect of sleep disruption on the ability to learn the correct location of the food reward (df=1, F=5.5972, p=0.0346. Figure 2, supp table 1). Specifically, fish in the sleep disturbed treatment performed on average significantly more incorrect choices before finding the food reward (5.65 incorrect choices) compared to fish in the non-disturbed sleep treatment (2.66 incorrect choices). As the days progressed, the mean number of incorrect choices by both treatment groups declined (Fig 2).

**Figure 2.**
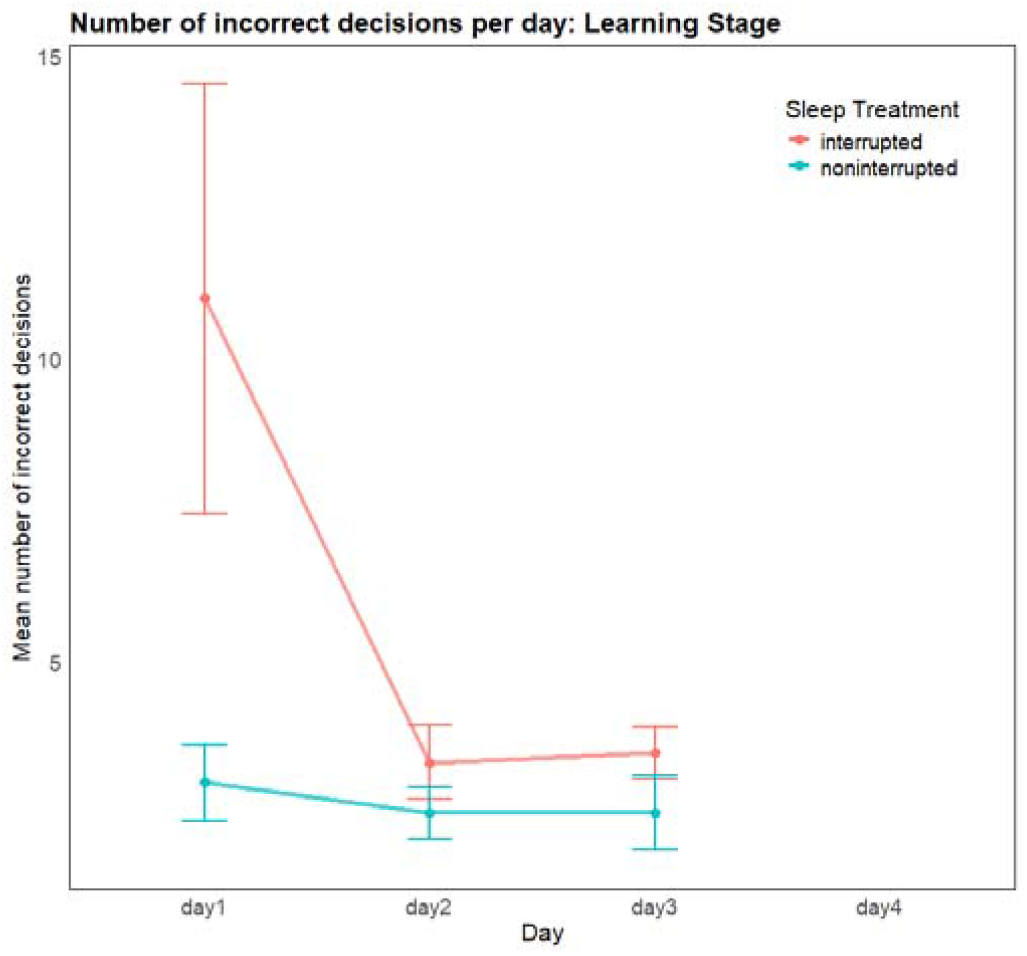
The mean number of incorrect decisions made by cleaner fish in two different sleep treatments (red: sleep disrupted; blue: sleep not disturbed) to find a food reward during the learning phase of the experiment. Mean incorrect choices per day; day 1: sleep disturbed fish 11, non-disturbed 3; day 2: sleep disturbed fish 3.3, non-disturbed 2.5; day 3: sleep disturbed fish 3.5, non-disturbed 2.5.

Similarly, we found a significant negative effect of sleep disruption on the time to find the correct food reward square (df=1, F=5.117, p=0.0472. Figure 3, supp table 2). Sleep disturbed fish took on average 794 seconds to find the food reward, while non-disturbed fish took 51.5 seconds. Again, fish in both groups improved over time and could find the food reward in a similar time by the third day.

**Figure 3.**
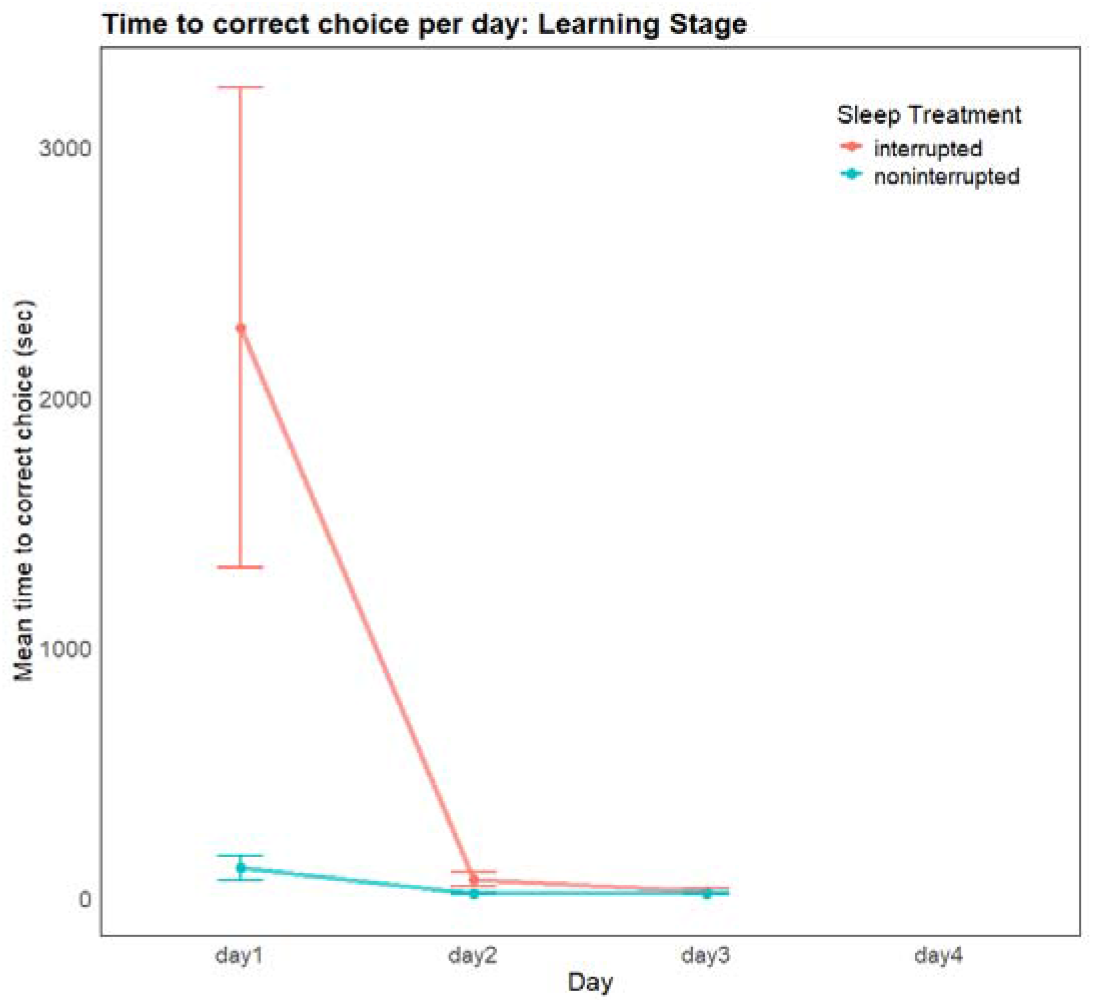
The mean time taken to find the food reward by cleaner fish in two different sleep treatments (red: sleep disrupted; blue: sleep not disturbed) during the learning phase of the experiment. Mean time per day; day 1: sleep disturbed fish 2278 sec, non-disturbed 120 sec; day 2: sleep disturbed fish 74.5 sec, non-disturbed 17.7 sec; day 3: sleep disturbed fish 29.7 sec, non-disturbed 16.8 sec.

### Impact of past sleep disruption on the ability to remember a cognitive task

We again found no significant difference in mean time to exit the door between the two treatments (df=1, F= 0.581, P=0.465).

In contrast to part of our second prediction, after the rest phase, we did not find a significant difference in the number of incorrect choices made before finding the food reward (df=1, F=1.937, p=0.197. Figure 4, supp table 3). Individuals that had been sleep disrupted during the previous learning phase performed on average 2.4 incorrect choices, while non-disturbed sleep individuals performed on average 1.5 incorrect choices.

**Figure 4.**
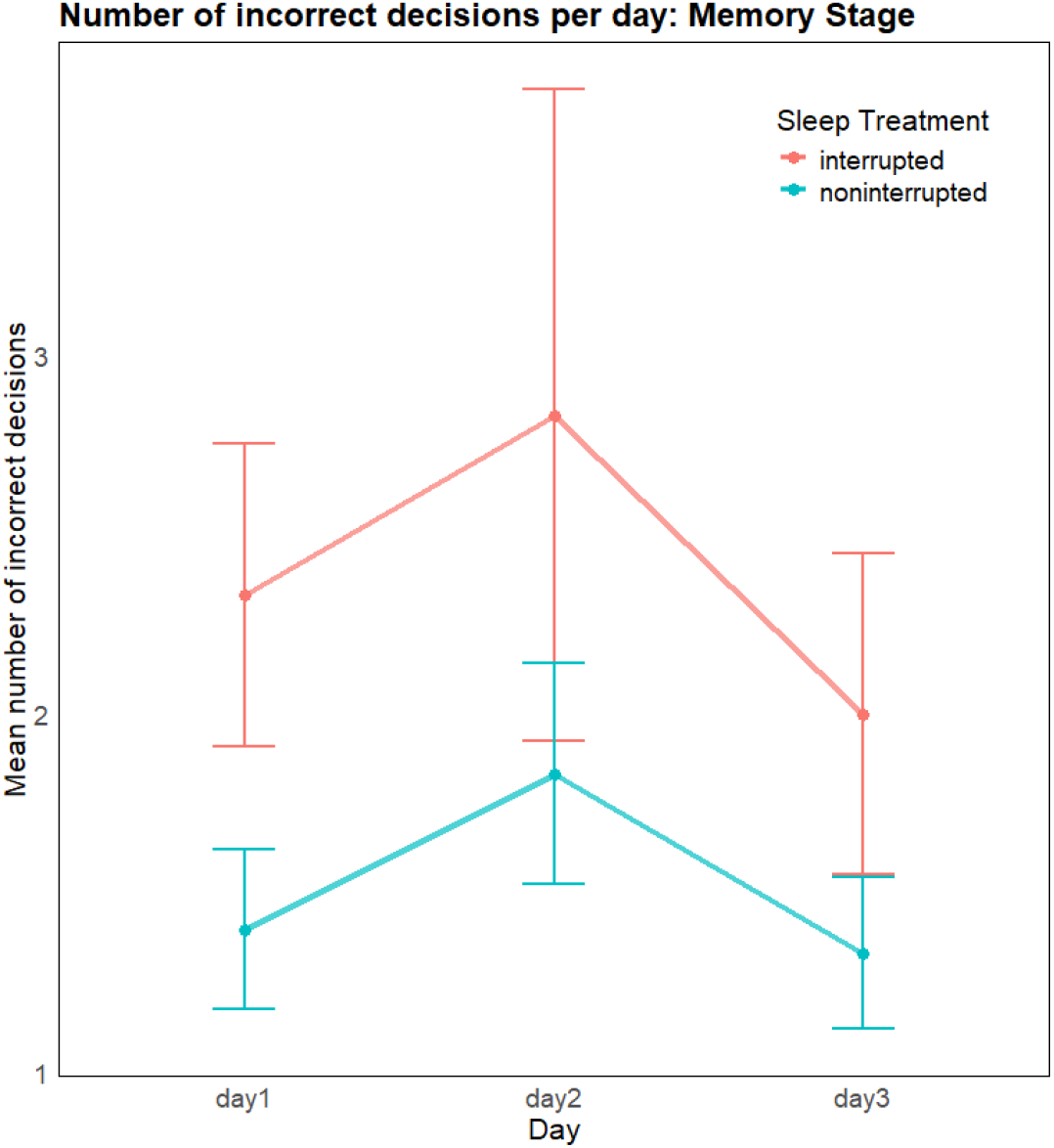
The mean number of incorrect decisions made by cleaner fish in two different sleep treatments (red: sleep disrupted; blue: sleep not disturbed) to find a food reward during the memory phase of the experiment. Mean incorrect choices per day; day 1: sleep disturbed fish 2.3, non-disturbed 1.4; day 2: sleep disturbed fish 2.8, non-disturbed 1.8; day 3: sleep disturbed fish 2, non-disturbed 1.3.

Likewise, we found no significant difference in the time to find the food reward (df=1, F=2.549, p=0.145. Figure 5, supp table 4). Individuals that had been sleep disrupted during the previous learning phase took on average 5.6 sec, whereas non-disturbed sleep individuals took on average 2.6 sec to find the food reward.

**Figure 5.**
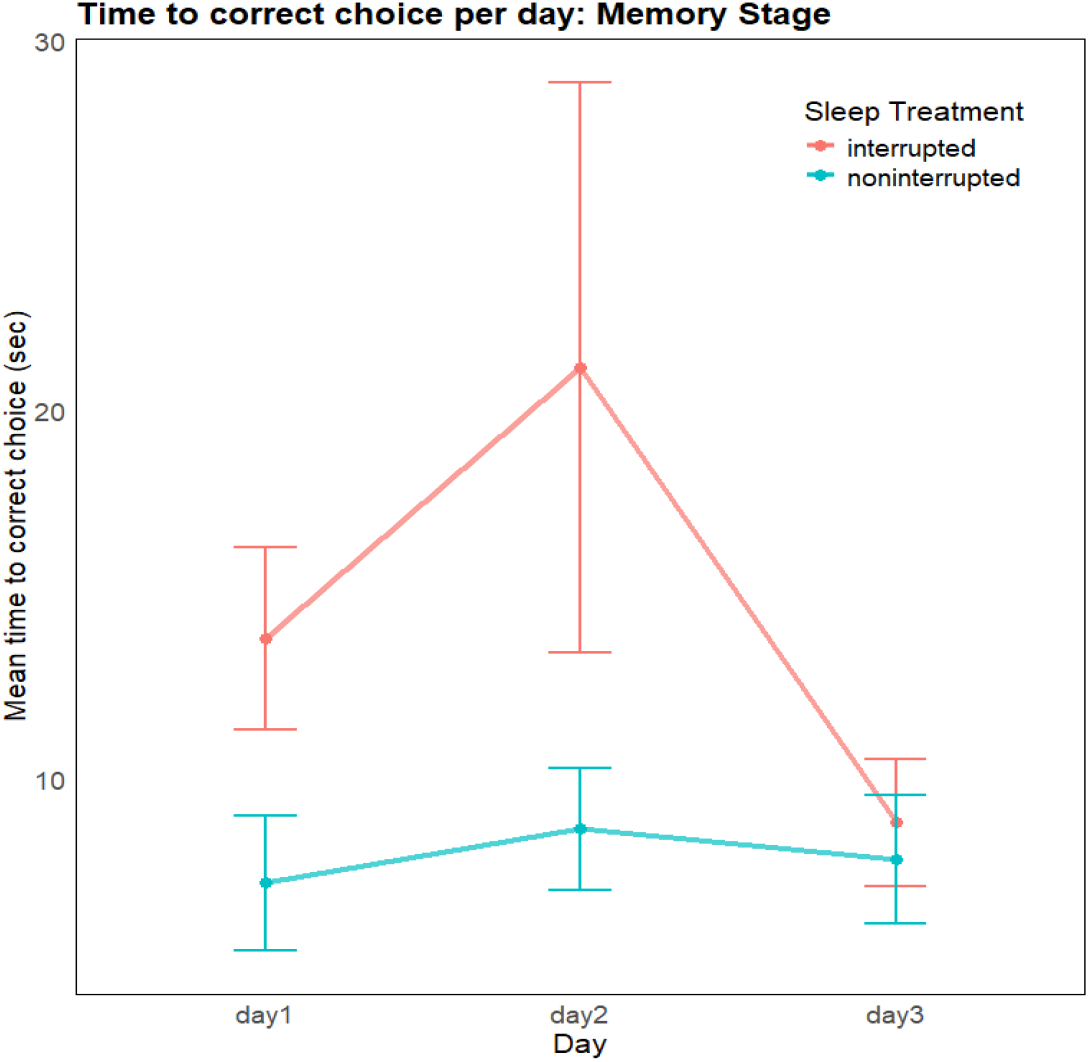
The mean time taken to find the food reward by cleaner fish in two different sleep treatments (red: sleep disrupted; blue: sleep not disturbed) during the memory phase of the experiment. Mean time per day; day 1: sleep disturbed fish 13.8 sec, non-disturbed 7.2 sec; day 2: sleep disturbed fish 21.2 sec, non-disturbed 8.8 sec; day 3: sleep disturbed fish 8.8 sec, non-disturbed 7.8 sec.

In both groups, individuals performed better at the food reward task in the memory phase compared to the learning phase. Generally, performances continued to get better as the memory phase progressed, however the pattern was not as clear as the continued improvement in completing the task in the learning phase.

## Discussion

We clearly demonstrate that disruptions to a normal sleep pattern can negatively affect how quickly and accurately an animal performs a novel cognitive task. We used as our model species the cleaner fish, to provide further evidence of an important function of sleep in non-mammalian and non-avian taxa. Despite retaining small mean differences overtime, we did not observe any significant sleep treatment differences in ability to perform the cognitive task during the memory phase of the experiment. Indeed, the ability to perform the task continued to improve in both treatment groups throughout both the learning phase and to a lesser extent the memory phase of the experiment. We demonstrate that any negative effects of sleep disruption could be most apparent when first encountering a new cognitive challenge. Nevertheless, our results support the claim that sleep plays an important role in mental processes, such as learning, across vertebrate taxa.

Sleep is predicted to maintain optimal brain functioning to support waking cognition (Alhola & Polo-Kantola 2007). The ability to learn is a critical function that allows strategic adaptation to changing environmental demands. Cleaner fish inhabit structurally complex physical environments and dynamic social environments and to be successful they require the ability to learn and to adapt their behavioural responses to changes accordingly. For example, it is estimated that wild cleaner fish remove up to 1200 ectoparasites each day (Grutter, 1996) from anywhere between 800 to 3000 client fish (Grutter, 1995; Triki et al. 2018; Wismer et al., 2014). In this context, cleaner fish must learn and remember information about their clients, including past positive or negative interactions, and identifying which clients are residents or visitors to their cleaning territories. We know that cleaner fish are also capable of social learning, with juvenile fish learning strategies to interact with clients by observing cleaner adults interacting with clients (Truskanov et al. 2020). We also know that cleaner fish are not the only fish species capable of learning, for instance archer fish (*Toxotes* sp.) learn complex skills from observing other group members (Schuster et al. 2006), while guppies are capable of learning and anticipating the path of a mechanical fish (Bierbach et al. 2022) and successfully completing reverse learning choice tasks (Buechel et al. 2018). It is therefore likely that other fish will also be negatively impacted by sleep disruption when learning novel tasks.

In cleaner fish, the negative effects of sleep disruption were most apparent at the beginning of the learning phase of the experiment. Our results may suggest differences in the timing of the performance of operant behaviours, i.e. those that that produce consequences and may be repeated if they provide a reward (Toates 2012), with sleep-interrupted fish taking longer to find the food reward on the first day after sleep disruption. An explanation for our finding could be that fish in the sleep-disrupted treatment are less active or motivated. Indeed, a similar finding has been observed in Australian magpies, where sleep loss impaired motivation and cognitive performance (Johnsson et al. 2022). In contrast, sleep disruption appeared to have less of an effect on reinforcement learning on subsequent days, with the time taken to find the food reward similar between both sleep treatments on days two and three of the experiment. Reinforcement learning influences behaviour via either a positive or negative feedback response and can be somewhat affected by sleep loss (Whitney et al. 2015; Gerhardsson et al. 2020). We recommend that future experiments investigating the association between sleep and learning should include a reverse learning element, which has shown in humans to amplify the impairment caused by sleep loss (Whitney et al. 2015).

We did not find a significant negative effect of sleep disruption on the ability of cleaner fish to recall the food reward cue after several days post learning phase. In humans, memory functions comprise three major subprocesses, i.e., encoding, consolidation, and retrieval (Straube 2012). Research has indicated that sleep after learning can have an important influence on memory consolidation. Rather we found that the performance of fish in both sleep treatments increased with time and exposure to the task in both the learning and to a lesser extent the memory stages of the experiment. Our findings do not rule out an important role for sleep in memory consolidation in fish, specifically non-declarative memory, but potentially represents a limitation in our experimental design. For example, the cognitive task may have been too simple or was repeated too often during the experimental period. Another possible explanation is that our periods of sleep disruption were too short to affect the consolidation and retrieval of memories associated with reinforcement learning. In our experiment fish were still allowed several hours sleep each night, albeit broken into discreet periods of time by the automated light. It is possible that a more complex cognitive task, including a reverse learning component, or longer periods of sleep deprivation each night may have increased cognitive and recall impairment.

The involvement of sleep in memory processing has been tested for at least a century (Rasch & Born 2013; Jenkins & Dallenbach 1924). Yet, sleep research is still overwhelmingly biased towards humans, other mammals, and laboratory model species (Toth & Bhargava 2013; Blumberg et al. 2020). Experimental evidence, has demonstrated behavioural, anatomical, genetic, and pharmacological conservation of sleep between fish (largely in zebrafish) and mammals, suggesting that research in fish can inform our understanding of mammalian sleep (Lee et al. 2022). Despite difficulties finding neuronal sleep signatures in fishes, due to the absence of a conventional neocortex, bony fishes do possess a homologue, the dorsal pallium, and neural signatures of sleep have been observed (Leung et al. 2019). Our evidence that sleep has an impact on mental processing in fish, in combination with the conservation of sleep and the presence of homologous structures in the vertebrate brain, suggests that sleep has played an important role in animal cognitive processes since early in the evolutionary history of vertebrate taxa.

Our study provides further evidence that sleep is functionally important in non-mammalian animals. Unlike previous studies on non-mammal species, we use an ecologically relevant stimulus to test learning ability and memory consolidation/retention using the cleaner fish. We found that the negative impacts of sleep disruption were most apparent initially, likely impacting cognition, activity levels and motivation, which in turn slowed the learning process. Our results do not uncover which aspects of cognition were most affected by sleep loss. However, we speculate that the association between sleep and mental processes is widespread in vertebrates and may have coevolved or facilitated the ability of animals to perform complex cognitive behaviours.

## Supplementary material

**Table 1.** Number of incorrect choices before food reward (Learning Phase)

|  | Df | Sum Sq | Mean Sq | F value | Pr(>F) |
| --- | --- | --- | --- | --- | --- |
| sleep_treatment | 1 | 77.66 | 77.66 | 5.972 | 0.0346 * |
| day | 1 | 0.68 | 0.68 | 0.052 | 0.8240 |
| Residuals | 10 | 130.04 | 13.00 |  |  |

**Table 2.** Time to correct choice (Learning Phase)

|  | Df | Sum Sq | Mean Sq | F value | Pr(>F) |
| --- | --- | --- | --- | --- | --- |
| sleep_treatment | 1 | 4958787 | 4958787 | 5.117 | 0.0472 * |
| day | 1 | 353 | 353 | 0.000 | 0.9852 |
| Residuals | 10 | 9690710 | 969071 |  |  |

**Table 3.** Number of incorrect choices before food reward (Memory Phase)

|  | Df | Sum Sq | Mean Sq | F value | Pr(>F) |
| --- | --- | --- | --- | --- | --- |
| sleep_treatment | 1 | 6.458 | 6.458 | 1.937 | 0.197 |
| day | 1 | 0.502 | 0.502 | 0.151 | 0.707 |
| Residuals | 9 | 30.011 | 3.335 |  |  |

**Table 4.** Time to correct choice (Memory Phase)

|  | Df | Sum Sq | Mean Sq | F value | Pr(>F) |
| --- | --- | --- | --- | --- | --- |
| sleep_treatment | 1 | 389.0 | 389.0 | 2.549 | 0.145 |
| day | 1 | 2.5 | 2.5 | 0.017 | 0.900 |
| Residuals | 9 | 1373.3 | 152.6 |  |  |

## References

Acerbi, A. & Nunn, C.L. 2011. Predation and the Phasing of Sleep: An Evolutionary Individual-Based Model. Animal Behaviour 81 (4): 801–11. 10.1016/j.anbehav.2011.01.015.

Anafi, R.C., Kayser, M.S. & Raizen, D.M. 2019. Exploring Phylogeny to Find the Function of Sleep. Nature Reviews Neuroscience 20 (2): 109–16. 10.1038/s41583-018-0098-9.

Arble, D.M., Bass, J., Behn, C.D., Butler, M.P., Challet, E., Czeisler, C., Depner, C.M. et al. 2015. Impact of Sleep and Circadian Disruption on Energy Balance and Diabetes: A Summary of Workshop Discussions. Sleep 38 (12): 1849–60. 10.5665/sleep.5226.

Backhaus, J., Junghanns, K., Born, J., Hohaus, K., Faasch, F., & Hohagen, F. (2006). Impaired declarative memory consolidation during sleep in patients with primary insomnia: influence of sleep architecture and nocturnal cortisol release. Biological psychiatry, 60(12), 1324–1330.

Beyaert, L., Greggers U., & Menzel, R. 2012. Honeybees Consolidate Navigation Memory during Sleep. Journal of Experimental Biology 215 (22): 3981–88. 10.1242/jeb.075499.

Bierbach D, Gómez-Nava L, Francisco FA, Lukas J, Musiolek L, Hafner VV, Landgraf T, Romanczuk P. & Krause J. 2022. Live fish learn to anticipate the movement of a fish-like robot. Bioinspir Biomim. 17(6).

Born, J., Rasch, B. & Gais, S. 2006. Sleep to Remember. The Neuroscientist: A Review Journal Bringing Neurobiology, Neurology and Psychiatry 12 (5): 410–24. 10.1177/1073858406292647.

Buechel S.D., Boussard A., Kotrschal A., van der Bijl W. & Kolm N. 2018. Brain size affects performance in a reversal-learning test. Proc Biol Sci. 31;285(1871):20172031

Campbell, S.S., & Tobler, I. 1984. Animal Sleep: A Review of Sleep Duration across Phylogeny. Neuroscience & Biobehavioral Reviews 8 (3): 269–300. 10.1016/0149-7634(84)90054-X.

Capellini, I., McNamara, P., Preston, B.T., Nunn, C.L. & Barton, R.A. 2009. Does Sleep Play a Role in Memory Consolidation? A Comparative Test. PLoS ONE 4 (2). 10.1371/journal.pone.0004609.

Cappuccio, F.P., D’Elia, L., Strazzullo, P. & Miller M.A. 2010. Sleep Duration and All-Cause Mortality: A Systematic Review and Meta-Analysis of Prospective Studies. Sleep 33 (5): 585–92. 10.1093/sleep/33.5.585.

Cirelli, C., & Tononi, G. 2008. Is Sleep Essential? PLOS Biology 6 (8): e216. 10.1371/journal.pbio.0060216.

Derégnaucourt, S., Mitra, P.P., Fehér, O., Pytte, C. & Tchernichovski, O. 2005. How Sleep Affects the Developmental Learning of Bird Song. Nature 433 (7027): 710–16. 10.1038/nature03275.

Diekelmann, S. & Born, J. 2010. The Memory Function of Sleep. Nature Reviews. Neuroscience 11 (2): 114–26. 10.1038/nrn2762.

Dinges, D.F., Pack, F., Williams, K., Gillen, K.A., Powell, J.W., Ott, G.E., Aptowicz, C. & Pack, A.I. 1997. Cumulative Sleepiness, Mood Disturbance, and Psychomotor Vigilance Performance Decrements During a Week of Sleep Restricted to 4–5 Hours per Night. Sleep 20 (4): 267–77. 10.1093/sleep/20.4.267.

Frank, M.G., Waldrop, R.H., Dumoulin, M., Aton, S. & Boal, J.G. 2012. A Preliminary Analysis of Sleep-Like States in the Cuttlefish Sepia Officinalis. PLOS ONE 7 (6): e38125. 10.1371/journal.pone.0038125.

Gerhardsson, A., Porada, D.K., Lundström, J.N., Axelsson, J., Schwarz, J. Does insufficient sleep affect how you learn from reward or punishment? Reinforcement learning after 2 nights of sleep restriction. J Sleep Res, 2020,. 10.1111/jsr.13236

Hairston, Ilana S., Milton T. M. Little, Michael D. Scanlon, Monique T. Barakat, Theo D. Palmer, Robert M. Sapolsky, and H. Craig Heller. 2005. “Sleep Restriction Suppresses Neurogenesis Induced by Hippocampus-Dependent Learning.” Journal of Neurophysiology 94 (6): 4224–33. 10.1152/jn.00218.2005.

Hartse, K.M. 1994. “Sleep in Insects and Nonmammalian Vertebrates.” In Principles and Practice of Sleep Medicine, edited by M. H. Kryger, T Roth, and W.C. Dement, 1:95–104. Philadelphia: Elsevier Saunders.

Huber, Reto, M. Felice Ghilardi, Marcello Massimini, and Giulio Tononi. 2004. “Local Sleep and Learning.” Nature 430 (6995): 78–81. 10.1038/nature02663.

Jackson, Claire, Brian J. McCabe, Alister U. Nicol, Amanda S. Grout, Malcolm W. Brown, and Gabriel Horn. 2008. “Dynamics of a Memory Trace: Effects of Sleep on Consolidation.” Current Biology 18 (6): 393–400. 10.1016/j.cub.2008.01.062.

Jenkins J, Dallenbach K. Obliviscence during sleep and waking. Am J Psychol 1924; 35: 605–612.

Johnsson RD, Connelly F, Gaviraghi Mussoi J, Vyssotski AL, Cain KE, Roth TC 2nd, Lesku JA. Sleep loss impairs cognitive performance and alters song output in Australian magpies. Sci Rep. 2022 Apr 22;12(1):6645. doi: 10.1038/s41598-022-10162-7. PMID: 35459249; PMCID: PMC9033856.

Joiner, William J. 2016. “Unraveling the Evolutionary Determinants of Sleep.” Current Biologyll: CB 26 (20): R1073–87. 10.1016/j.cub.2016.08.068.

Kayser, Matthew S., Zhifeng Yue, and Amita Sehgal. 2014. “A Critical Period of Sleep for Development of Courtship Circuitry and Behavior in Drosophila.” Science 344 (6181): 269–74. 10.1126/science.1250553.

Keene, Alex C., and Erik R. Duboue. 2018. “The Origins and Evolution of Sleep.” Journal of Experimental Biology 221 (11). 10.1242/jeb.159533.

Krishnan, Harini C., Eric J. Noakes, and Lisa C. Lyons. 2016. “Chronic Sleep Deprivation Differentially Affects Short and Long-Term Operant Memory in Aplysia.” Neurobiology of Learning and Memory 134 (October): 349–59. 10.1016/j.nlm.2016.08.013.

Lee, D. A., Oikonomou, G., & Prober, D. A. (2022). Large-scale analysis of sleep in Zebrafish. Bio-protocol, 12(3), e4313–e4313.

Lee Kavanau, J. 1996. “Memory, Sleep, and Dynamic Stabilization of Neural Circuitry: Evolutionary Perspectives.” Neuroscience & Biobehavioral Reviews 20 (2): 289–311. 10.1016/0149-7634(95)00019-4.

Lesku, John A., and Niels C. Rattenborg. 2014. “Avian Sleep.” Current Biology 24 (1): R12–14. 10.1016/j.cub.2013.10.005.

Leung, Louis C., Gordon X. Wang, Romain Madelaine, Gemini Skariah, Koichi Kawakami, Karl Deisseroth, Alexander E. Urban, and Philippe Mourrain. 2019. “Neural Signatures of Sleep in Zebrafish.” Nature 571 (7764): 198–204. 10.1038/s41586-019-1336-7.

Lima, Steven L. 1998. “Stress and Decision Making under the Risk of Predation: Recent Developments from Behavioral, Reproductive, and Ecological Perspectives.” In Advances in the Study of Behavior, edited by Anders Pape Møller, Manfred Milinski, and Peter J. B. Slater, 27:215–90. Stress and Behavior. Academic Press. 10.1016/S0065-3454(08)60366-6.

Mateo, Jill M., and Robert E. Johnston. 2000. “Retention of Social Recognition after Hibernation in Belding’s Ground Squirrels.” Animal Behaviour 59 (3): 491–99. 10.1006/anbe.1999.1363.

Mazzotti, Diego Robles, Camila Guindalini, Walter André dos Santos Moraes, Monica Levy Andersen, Maysa Seabra Cendoroglo, Luiz Roberto Ramos, and Sergio Tufik. 2014. “Human Longevity Is Associated with Regular Sleep Patterns, Maintenance of Slow Wave Sleep, and Favorable Lipid Profile.” Frontiers in Aging Neuroscience 6. 10.3389/fnagi.2014.00134.

Mignot, Emmanuel. 2008. “Why We Sleep: The Temporal Organization of Recovery.” PLOS Biology 6 (4): e106. 10.1371/journal.pbio.0060106.

Nath, Ravi D., Claire N. Bedbrook, Michael J. Abrams, Ty Basinger, Justin S. Bois, David A. Prober, Paul W. Sternberg, Viviana Gradinaru, and Lea Goentoro. 2017. “The Jellyfish Cassiopea Exhibits a Sleep-like State.” Current Biology 27 (19): 2984-2990.e3. 10.1016/j.cub.2017.08.014.

Peigneux, Philippe, Steven Laureys, Xavier Delbeuck, and Pierre Maquet. 2001. “Sleeping Brain, Learning Brain. The Role of Sleep for Memory Systems.” NeuroReport 12 (18): A111.

Pinheiro-da-Silva, Jaquelinne, Steven Tran, and Ana Carolina Luchiari. 2018. “Sleep Deprivation Impairs Cognitive Performance in Zebrafish: A Matter of Fact?” Behavioural Processes 157 (December): 656–63. 10.1016/j.beproc.2018.04.004.

Portavella, M, J. P Vargas, B Torres, and C Salas. 2002. “The Effects of Telencephalic Pallial Lesions on Spatial, Temporal, and Emotional Learning in Goldfish.” Brain Research Bulletin 57 (3): 397–99. 10.1016/S0361-9230(01)00699-2.

Preston, Brian T., Isabella Capellini, Patrick McNamara, Robert A. Barton, and Charles L. Nunn. 2009. “Parasite Resistance and the Adaptive Significance of Sleep.” BMC Evolutionary Biology 9 (1): 7. 10.1186/1471-2148-9-7.

Rasch B, Born J. About sleep’s role in memory. Physiol Rev. 2013 Apr;93(2):681–766. doi: 10.1152/physrev.00032.2012. PMID: 23589831; PMCID: PMC3768102.

Rattenborg, Niels C., Horacio O. de la Iglesia, Bart Kempenaers, John A. Lesku, Peter Meerlo, and Madeleine F. Scriba. 2017. “Sleep Research Goes Wild: New Methods and Approaches to Investigate the Ecology, Evolution and Functions of Sleep.” Philosophical Transactions of the Royal Society B: Biological Sciences 372 (1734). 10.1098/rstb.2016.0251.

Rawashdeh, Oliver, Nancy Hernandez de Borsetti, Gregg Roman, and Gregory M. Cahill. 2007. “Melatonin Suppresses Nighttime Memory Formation in Zebrafish.” Science 318 (5853): 1144–46. 10.1126/science.1148564.

Rechtschaffen, Allan, and Bernard M. Bergmann. 2002. “Sleep Deprivation in the Rat: An Update of the 1989 Paper.” Sleep 25 (1): 18–24. 10.1093/sleep/25.1.18.

Ribeiro, Sidarta, Damien Gervasoni, Ernesto S. Soares, Yi Zhou, Shih-Chieh Lin, Janaina Pantoja, Michael Lavine, and Miguel A. L. Nicolelis. 2004. “Long-Lasting Novelty-Induced Neuronal Reverberation during Slow-Wave Sleep in Multiple Forebrain Areas.” PLoS Biology 2 (1): E24. 10.1371/journal.pbio.0020024.

Romer, Alfred Sherwood. 1959. The Vertebrate Story. University of Chicago Press.

Roth, Timothy C., Niels C. Rattenborg, and Vladimir V. Pravosudov. 2010. “The Ecological Relevance of Sleep: The Trade-off between Sleep, Memory and Energy Conservation.” Philosophical Transactions of the Royal Society B: Biological Sciences 365 (1542): 945–59. 10.1098/rstb.2009.0209.

Sauer, S., M. Kinkelin, E. Herrmann, and W. Kaiser. 2003. “The Dynamics of Sleep-like Behaviour in Honey Bees.” Journal of Comparative Physiology A 189 (8): 599–607. 10.1007/s00359-003-0436-9.

Schuster S, Wöhl S, Griebsch M, Klostermeier I. Animal cognition: how archer fish learn to down rapidly moving targets. Curr Biol. 2006 Feb 21;16(4):378–83. doi: 10.1016/j.cub.2005.12.037. PMID: 16488871.

Scullin, Michael K., and Donald L. Bliwise. 2015. “Is Cognitive Aging Associated with Levels of REM Sleep or Slow Wave Sleep?” Sleep 38 (3): 335–36. 10.5665/sleep.4482.

Shaw, P. J., C. Cirelli, R. J. Greenspan, and G. Tononi. 2000. “Correlates of Sleep and Waking in Drosophila Melanogaster.” Science (New York, N.Y.) 287 (5459): 1834–37. 10.1126/science.287.5459.1834.

Shaw, Paul J., Giulio Tononi, Ralph J. Greenspan, and Donald F. Robinson. 2002. “Stress Response Genes Protect against Lethal Effects of Sleep Deprivation in Drosophila.” Nature 417 (6886): 287–91. 10.1038/417287a.

Sherry, David F., and Jennifer S. Hoshooley. 2010. “Seasonal Hippocampal Plasticity in Food-Storing Birds.” Philosophical Transactions of the Royal Society B: Biological Sciences 365 (1542): 933–43. 10.1098/rstb.2009.0220.

Shettleworth, Sara J. 1995. “Comparative Studies of Memory in Food Storing Birds: From the Field to the Skinner Box.” In Behavioural Brain Research in Naturalistic and Semi-Naturalistic Settings, edited by E Alleva, A Fasolo, Hans-Peter Lipp, L Nadel, and L Ricceri, 159–92. The Netherlands: Kluwer Academic.

Shettleworth, Sara J.. 1998. Cognition, Evolution, and Behavior. 2nd ed. Oxford University Press.

Siegel, Jerome M. 2001. “The REM Sleep-Memory Consolidation Hypothesis.” Science 294 (5544): 1058–63. 10.1126/science.1063049.

Smith, C. 2001. “Sleep States and Memory Processes in Humans: Procedural versus Declarative Memory Systems.” Sleep Medicine Reviews 5 (6): 491–506. 10.1053/smrv.2001.0164.

Smith, C., and G. Kelly. 1988. “Paradoxical Sleep Deprivation Applied Two Days after End of Training Retards Learning.” Physiology & Behavior 43 (2): 213–16. 10.1016/0031-9384(88)90240-5.

Stickgold, Robert. 2006. “A Memory Boost While You Sleep.” Nature 444 (7119): 559–60. 10.1038/nature05309.

Stickgold, Robert, and Matthew P. Walker. 2005. “Memory Consolidation and Reconsolidation: What Is the Role of Sleep?” Trends in Neurosciences 28 (8): 408–15. 10.1016/j.tins.2005.06.004.

Straube, B. An overview of the neuro-cognitive processes involved in the encoding, consolidation, and retrieval of true and false memories. Behav Brain Funct 8, 35 (2012). 10.1186/1744-9081-8-35

Toates, F. (2012). Operant Behavior. In: Seel, N.M. (eds) Encyclopedia of the Sciences of Learning. Springer, Boston, MA. 10.1007/978-1-4419-1428-6_992

Tobler, I. 2011. “Phylogeny of Sleep Regulation.” In Principles and Practice of Sleep Medicine, edited by M. H. Kryger, T Roth, and W.C. Dement, 5th ed., 112–25. Philadelphia: Elsevier Saunders.

Tobler, Irene. 1995. “Is Sleep Fundamentally Different between Mammalian Species?” Behavioural Brain Research, The Function of Sleep, 69 (1): 35–41. 10.1016/0166-4328(95)00025-O.

Tobler, Irene, and Alexander A. Borbély. 1985. “Effect of Rest Deprivation on Motor Activity of Fish.” Journal of Comparative Physiology A 157 (6): 817–22. 10.1007/BF01350078.

Tononi, Giulio, and Chiara Cirelli. 2006. “Sleep Function and Synaptic Homeostasis.” Sleep Medicine Reviews 10 (1): 49–62. 10.1016/j.smrv.2005.05.002.

Vargas, Juan Pedro, Verner Peter Bingman, Manuel Portavella, and Juan Carlos López. 2006. “Telencephalon and Geometric Space in Goldfish.” European Journal of Neuroscience 24 (10): 2870–78. 10.1111/j.1460-9568.2006.05174.x.

Vertes, Robert P. 2004. “Memory Consolidation in Sleep: Dream or Reality.” Neuron 44 (1): 135–48. 10.1016/j.neuron.2004.08.034.

Vorster, Albrecht P., and Jan Born. 2015. “Sleep and Memory in Mammals, Birds and Invertebrates.” Neuroscience and Biobehavioral Reviews 50 (March): 103–19. 10.1016/j.neubiorev.2014.09.020.

Wagner, Ullrich, Steffen Gais, Hilde Haider, Rolf Verleger, and Jan Born. 2004. “Sleep Inspires Insight.” Nature 427 (6972): 352–55. 10.1038/nature02223.

Walker MP. The role of slow wave sleep in memory processing. J Clin Sleep Med. 2009 Apr 15;5(2 Suppl):S20–6. PMID: 19998871; PMCID: PMC2824214.

Walker, MP., and R. Stickgold. 2004a. “Sleep-Dependent Learning and Memory Consolidation.” Neuron 44 (1): 121–33. 10.1016/j.neuron.2004.08.031.

Walker, MP., and R. Stickgold. 2004b. “Sleep-Dependent Learning and Memory Consolidation.” Neuron 44 (1): 121–33. 10.1016/j.neuron.2004.08.031.

Walker, MP., and R. Stickgold. 2006. “Sleep, Memory, and Plasticity.” Annual Review of Psychology 57: 139–66. 10.1146/annurev.psych.56.091103.070307.

Whitney, P., Hinson, J. M., Jackson, M. L., & Van Dongen, H. P. A. (2015). Feedback blunting: Total sleep deprivation impairs decision making that requires updating based on feedback. Sleep, 38, 745–754.

Sharon Wismer, Ana I Pinto, Zegni Triki, Alexandra S Grutter, Dominique G Roche, Redouan Bshary 2019. Cue-based decision rules of cleaner fish in a biological market task. Animal behaviour 158: 249–260

Worley, Susan L. 2018. “The Extraordinary Importance of Sleep.” Pharmacy and Therapeutics 43 (12): 758–63.

Yokogawa, Tohei, Wilfredo Marin, Juliette Faraco, Guillaume Pézeron, Lior Appelbaum, Jian Zhang, Frédéric Rosa, Philippe Mourrain, and Emmanuel Mignot. 2007. “Characterization of Sleep in Zebrafish and Insomnia in Hypocretin Receptor Mutants.” PLoS Biology 5 (10). 10.1371/journal.pbio.0050277.

Zimmerman, John E., Nirinjini Naidoo, David M. Raizen, and Allan I. Pack. 2008. “Conservation of Sleep: Insights from Non-Mammalian Model Systems.” Trends in Neurosciences 31 (7): 371–76. 10.1016/j.tins.2008.05.001.

Yuanyuan Xie and Richard I. Dorsky (2017) Development of the hypothalamus: conservation, modification and innovation. 2017. Development. 144(9): 1588–1599. doi: 10.1242/dev.139055

